# Agentic AI Integrated with Scientific Knowledge: Laboratory Validation in Systems Biology

**DOI:** 10.1101/2025.06.24.661378

**Authors:** Daniel Brunnsåker, Alexander H. Gower, Prajakta Naval, Erik Y. Bjurström, Filip Kronström, Ievgeniia A. Tiukova, Ross D. King

**Affiliations:** Department of Computer Science and Engineering, Chalmers University of Technology and University of Gothenburg, SE-41296 Gothenburg, Sweden; Department of Life Sciences, Chalmers University of Technology, SE-41296 Gothenburg, Sweden; Division of Industrial Biotechnology, KTH Royal Institute of Technology, SE-10691 Stockholm, Sweden; Department of Chemical Engineering and Biotechnology, University of Cambridge, Cambridge CB3 0AS, United Kingdom; The Alan Turing Institute, London NW1 2DB, United Kingdom

**Keywords:** Laboratory Automation, Systems Biology, Machine Learning, Inductive Logic Programming, Automation of Science, Large Language Models

## Abstract

Automation is transforming scientific discovery by enabling systematic exploration of complex hypotheses. Large language models (LLMs) perform well across diverse tasks and promise to accelerate research, but often struggle to interact with logical structures. Here we present a framework integrating LLM-based agents with laboratory automation, guided by a logical scaffold incorporating symbolic relational learning, structured vocabularies, and experimental constraints. This integration reduces output incoherence and improves reliability in automated workflows.

We couple this AI-driven approach to automated cell-culture and metabolomics platforms, enabling hypothesis validation and refinement, yielding a flexible system for scientific discovery.

We validate the system in *Saccharomyces cerevisiae*, identifying novel interactions, including glutamate-induced synergistic growth inhibition in spermine-treated cells and aminoadipate’s partial rescue of formic-acid stress. All hypotheses, experiments, and data are captured in a graph database employing controlled vocabularies. Existing ontologies are extended, and a novel representation of scientific hypotheses is presented using description logics. This work enables a more reliable, machine-driven discovery process in systems biology.

## Introduction

Artificial intelligence (AI) and laboratory automation are rapidly transforming science. AI enables the analysis of data that would otherwise be intractable and, when integrated with laboratory robotics, can automate scientific research— a capability essential for addressing the increasingly complex problems found in modern science (1–3). For example, comparatively simple model organisms, such as *Saccharomyces cerevisiae* (baker’s yeast) and *Escherichia coli*, exhibit complexity far beyond what humans can interpret or validate experimentally within a reasonable timeframe (4). Reliable, scalable scientific discovery under these conditions requires integrated automated experimentation and computational analysis (4, 5).

The efficacy of scientists is determined by their scientific ideas, agency (the tools and techniques available to them), and collaborative capacity. For AI scientists, deciding what agency we grant them is an important design question. In particular, real-world agency—the ability to directly control experiments—is vital for scientific discovery. Integration with laboratory robotics was a focus of the previous generation of robot scientists; these model-driven systems generated hypotheses in tightly controlled experimental spaces and collected empirical data in only a few data modalities (4, 6–8). Several variations of this approach has been applied in fields such as materials development and chemistry (9, 10). Yet the next generation of robot scientists must contend with increasingly complex scientific questions that demand more powerful tools for hypothesis generation, multimodal data integration, experimental design, and analysis (1).

One promising direction is to grant robot scientists access to large language models (LLMs), which have enabled AI systems to achieve impressive results in tasks that previously required human intelligence, including scientific discovery (3, 11). LLM-augmented approaches offer adaptability in how problems are formulated and in expanding the experimental scope—a major advantage in the natural sciences. Consequently, we see increasingly widespread use and impact of LLMs in scientific discovery, often via orchestrated multi-agent systems (12–19).

The adaptability of LLM-augmented approaches is particularly beneficial in systems biology research, which involves complex interaction networks and vast spaces of experimental conditions and data modalities (20). However, LLM outputs are often inconsistent, difficult to verify, and rarely subjected to rigorous experimental evaluation—issues compounded by siloed data practices that limit transparency and reuse (18). A further weakness of LLMs remains their inability, or at best inefficiency and unreliability, when interacting with formal languages and logical structures. Recent agentic systems mitigate this issue by granting the LLM agency to write and execute programs in high-level languages such as Python (17). However, these approaches generally rely on loosely typed message passing, shallow artifact tracking, and lack unified domain representations. As a result, they can generate brittle, unvalidated outputs that are difficult to systematically audit (18).

Agentic systems for scientific discovery should make the best use of available scientific data and models for ideation, and report in a way that enables good collaboration and recording of knowledge. In many cases, including in systems biology, there is a wealth of structured data encoded according to well-defined rules. There is thus a need for frameworks that retain the flexibility of LLMs while grounding them in formal logic, exploiting background knowledge and meeting transparent data-sharing requirements (12, 17–19, 21).

Here we introduce a hybrid approach that couples language-driven flexibility with inductive logic and formal ontologies (see Fig. 1). The framework combines LLMs with the precision of relational learning (inductive logic programming, ILP) and community-adopted scientific ontologies to generate hypotheses grounded in empirical data and structured biological knowledge. The hypotheses are prioritised based on prior experimental results and can, thanks to their structured format, be automatically tested on laboratory automation platforms such as robotic liquid handlers, automated cultivation systems, and mass-spectrometry–based metabolomics (8, 22). All metadata, logs, and results can then be stored in a graph database that ensures transparency, reproducibility, and efficient reuse.

**Fig. 1.**
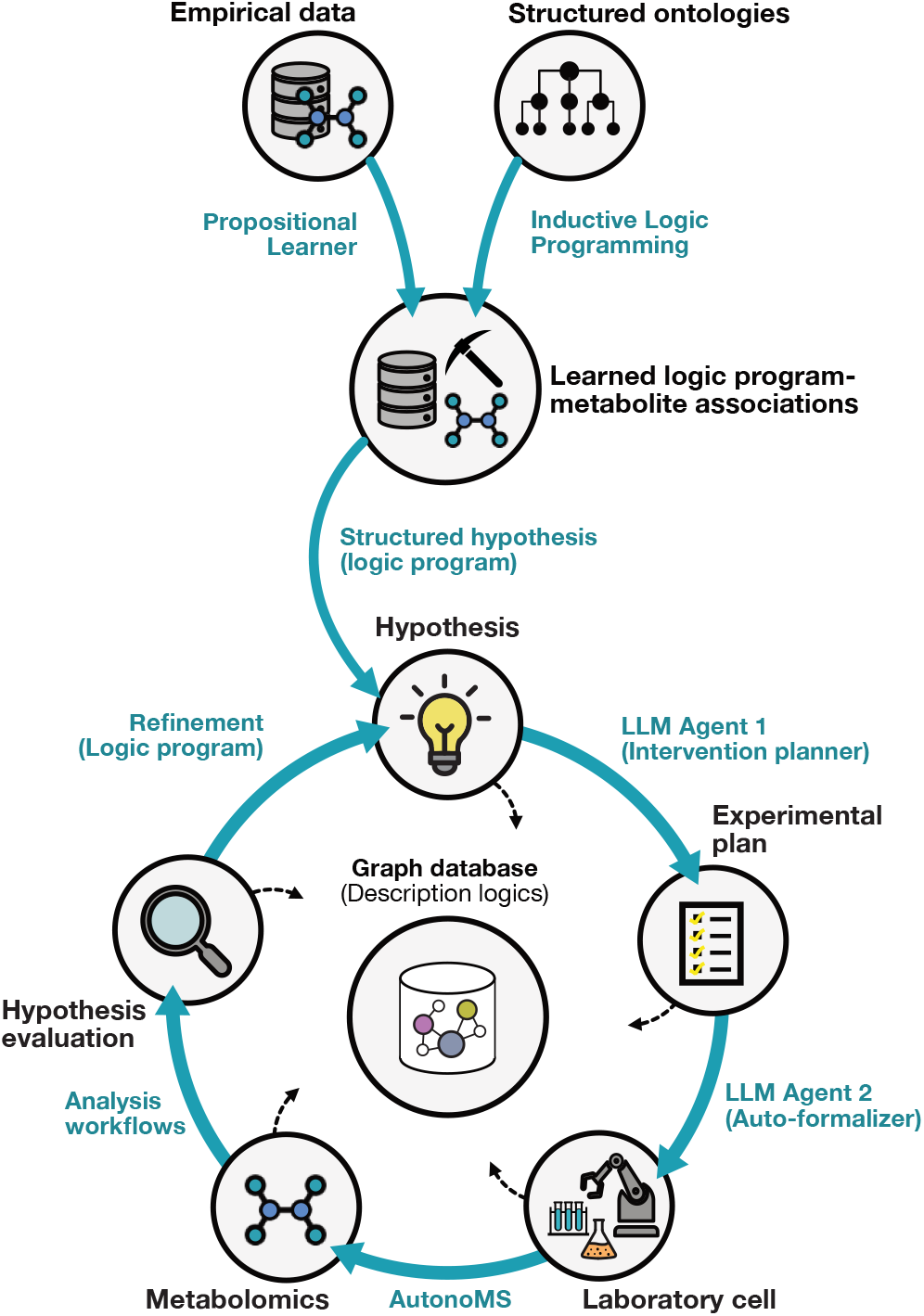
An automated experimental framework enabling end-to-end biological discovery—from hypothesis generation to data integration. Hypotheses are generated by mining patterns from a Datalog database containing structured facts about yeast phenotype, physiology, and metabolism, using inductive logic programming. These are linked to metabolomics data via regression. Interventions to test each hypothesis are designed by a large language model (LLM), which incorporates real-world experimental constraints. The interventions are auto-formalized into machine-readable protocols and executed by an automated laboratory cell. Time-series growth data are collected throughout the experiments, followed by automated sample processing and acquisition of endpoint metabolic profiles via ion-mobility mass spectrometry, mediated by AutonoMS (22). All data—including results, intermediate outputs, and metadata are curated, analyzed, and integrated into a graph database for downstream access and reuse.

By integrating symbolic learning with ontologies and statistical inference, the framework overcomes key limitations of standard agentic LLM systems: it improves traceability, reduces speculative outputs, and enables higher degrees of auditability across the entire discovery loop.

This framework was evaluated through automated experimental testing of several data-driven hypotheses. For example, we observed glutamate-induced synergistic growth inhibition in spermine-treated cells and arginine-mediated enhancement of caffeine toxicity in caffeine-treated cells. We further applied it to test a metabolomics-informed prediction that led to the discovery of aminoadipate-mediated rescue of formic acid stress. These results demonstrate the framework’s potential for automated, scalable, reproducible discovery in biological systems.

## Results

### Automated Systems Biology Research

We built an automated experimental and computational framework by integrating LLMs; structured community databases; robotic liquid handling stations; an automated cell-culturing and sampling platform; an ion mobility–based mass spectrometry metabolomics workflow; and computational tools in R, Python and Prolog (8, 22–26). See Fig. 1 for a graphical overview of the process.

This framework is designed to automatically generate and experimentally evaluate hypotheses about cell biology. Testable hypotheses are generated by (i) data-mining semantically meaningful facts about *S. cerevisiae* phenotype and metabolism—extracted from structured community databases (27–31)—(ii) applying pattern mining to generate a logical rule (25, 26), and (iii) applying machine learning on metabolomics data to learn a testable implication associated to the rule (32). The generated hypotheses are filtered and prioritized using prior experimental data. LLM agents then design experimental interventions to test these hypotheses and further formalize them into a machine-readable format with executable real-world constraints (33). Automated experiments are executed for each hypothesis, producing and processing biological samples to generate time-series growth data and endpoint metabolic profiles via ion-mobility-based mass spectrometry. After each run, the data are automatically curated and analysed, and presented in a concise, user-readable summary.

Each step of the process and accompanying experimental results are recorded in a graph database, including all LLM queries and responses, to ensure transparency and facilitate efficient reuse of data. Experimental data and metadata are saved using controlled vocabularies, utilising an extended version of the Ascomycete Phenotype Ontology (APO) (27, 28). Hypotheses and their implications are represented using description logics. This approach allows reusing existing data when a generated hypothesis matches a previously executed experiment, as demonstrated for one hypothesis below (see *hypothesis 9* in table 1).

**Table 1.** Summary of hypotheses selected for testing. The evaluation shows, for each hypothesis, whether the change in intracellular accumulation and the effect on resistance were (i) observed and (ii) in the predicted direction, and whether the negative control showed specificity. “-”, “+” indicates the observed or predicted sign of the change. “~” denotes an inconclusive change in accumulation. *\*Iterative hypothesis generated through data generated in hypothesis 4*.. *\*\*Hypothesis tested using already acquired data. †Experiment tests inverse hypothesis, i*.*e. nutrient supplementation rather than reduction, and reversed effect*.

| Generated Hypothesis | Change in accumulation |  | Effect on resistance |  | Intervention specificity |
| --- | --- | --- | --- | --- | --- |
|  | Observed | Predicted | Observed | Predicted |  |
| *Hypothesis 1* ↓ Arginine ⇒ ↑ Caffeine resistance† | + | + | – | – | Yes |
| *Hypothesis 2* ↑ Glutamate ⇒ ↓ Spermine resistance | ~ | + | – | – | Yes |
| *Hypothesis 3* ↑ Glutamate ⇒ ↓ Formic acid resistance | ~ | + | – | + | No |
| *Hypothesis 4* ↓ Glutamine ⇒ ↓ Acetic acid resistance† | + | + | – | + | No |
| *Hypothesis 5* ↑ Lysine ⇒ ↓ Hyperosmotic stress resistance (at 30% (w/v) sucrose) | + | + | + | – | Yes |
| *Hypothesis 6* ↓ Arginine ⇒ ↓ Lithium(+1) resistance (at 0.1M lithium chloride)† | + | + | ~ | + | No |
| *Hypothesis 7* ↑ Proline ⇒ ↑ Lactic acid resistance | + | + | ~ | + | No |
| *Hypothesis 8\** ↑ Amino adipate ⇒ ↑ Formic acid resistance (at 10mM formic acid) | – | + | + | + | Yes |
| *Hypothesis 9\*\** ↑ Proline ⇒ ↓ Formic acid resistance | ~ | + | ~ | – | No |

### Linking metabolites to traits for hypothesis discovery

To generate testable, data-driven hypotheses based on prior scientific knowledge, we constructed a Datalog database of approximately 60,000 phenotypical, physiological, and metabolic relations for *S. cerevisiae*. This database integrates curated datasets from several different sources, structured by established ontologies, and encoded as logic facts to enable structured reasoning (28–30). Datalog’s declarative structure enables clear expression of biological relationships and efficient querying across complex relational datasets (34).

To identify patterns that could explain observed phenotypes, we applied ILP, a methodology well-suited to learning interpretable, well-defined logic programs (i.e., rules) from structured, relational data (25, 26, 35). ILP-based pattern mining on this structured database yielded 735 distinct logic programs, 390 of which mapped uniquely to positive examples (gene deletion strains) from the metabolomics dataset of Mülleder et al. (32). We propositionalised each program into a binary feature (denoting whether the program covers a given example) and fitted a linear regression model to link these logic programs to metabolite measurements through regression coefficients. This produced 1933 independent hypotheses across 16 proteinogenic amino acids (see Fig. 2).

**Fig. 2.**
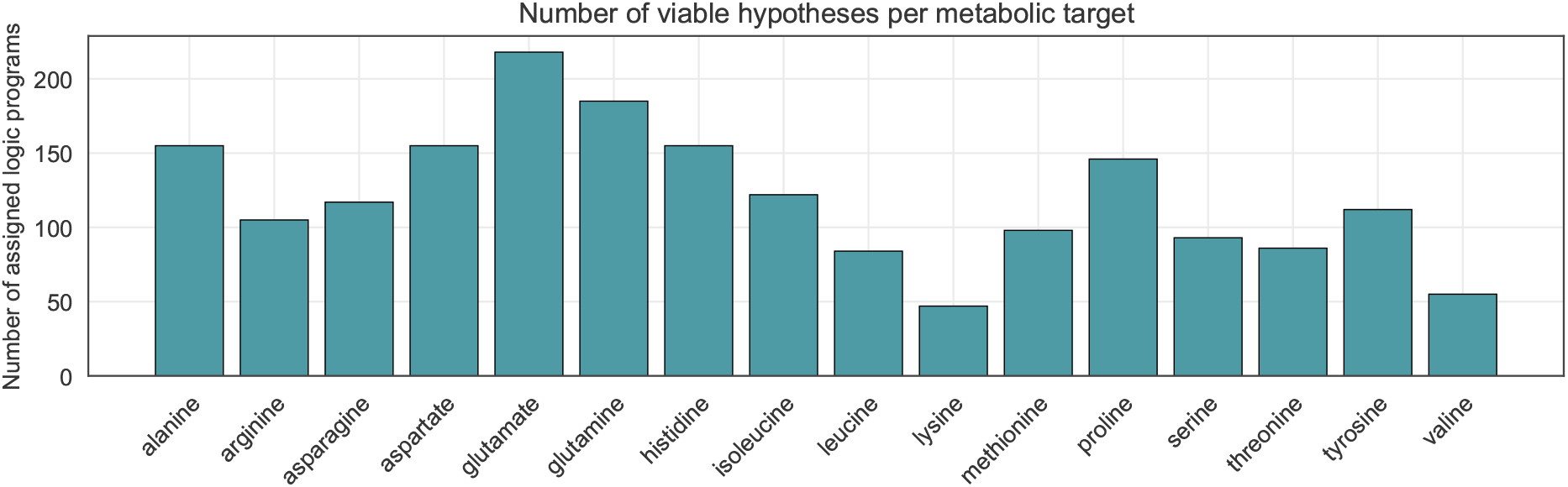
Viable hypothesis counts per amino acid. Bar chart showing, for each amino acid, the number of logic programs with non-zero regression coefficients—i.e., the total number of viable hypotheses generated for that amino acid.

An overview of the hypothesis space can be seen in Sup. Note 1 and Sup. Table 1. A logic program and learned association can be seen below (also see *hypothesis 6* in table 1).

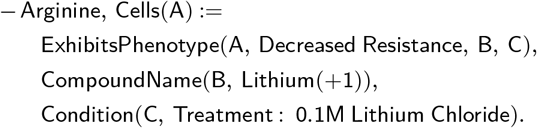

In this context, this program can be interpreted as: cells (A) which have decreased resistance to lithium (B) when treated with 0.1M lithium chloride (C) are associated with lower intracellular arginine levels. The link between arginine levels and the resistance phenotype suggests the testable hypothesis that arginine supplementation will restore lithium resistance (i.e. ↑ Arginine =⇒ ↑ Lithium(+1) resistance).

### Automated experiments identify interaction effects

To evaluate the utility of the framework we conducted automated investigations on chemical stressor and nutrient intervention interactions (the most common type of hypothesis generated in the previous step). For each amino acid, the framework selected the highest-weighted hypothesis, chose an appropriate negative control from the model’s learned associations (see Methods), and ran the corresponding experiment—producing time-series growth curves, endpoint metabolic profiles, effect sizes, and statistical evaluations.

Investigations were performed on five proteinogenic amino acids: L-glutamate, L-arginine, L-proline, Lglutamine, and L-lysine. Most investigations revealed significant interaction effects on growth in stressor-treated cells upon amino acid supplementation, several of which have not been previously discovered. An overview of all investigations is presented in table 1, with full statistical details in Sup. Note 7 and Sup. Table 2.

*Hypothesis 1* (see table 1) was consistent with acquired empirical data (shown in Fig. 3A). The addition of L-arginine alone caused a modest reduction in AUC (area under the growth curve) (−2.5% per mM, *p <* 0.05), whereas caffeine treatment showed an inhibitory effect (−53.4%, *p <* 0.001). Their combined application resulted in a synergistic decrease relative to independent effects (−19.0% per mM, *p <* 0.001), while the L-alanine negative control yielded a smaller decrease (−31.6% at 5mM, *p <* 0.01), confirming intervention specificity. Intracellular L-arginine accumulation increased significantly under high extracellular L-arginine supplementation (*p <* 0.01), confirming the validity of the intervention.

**Fig. 3.**
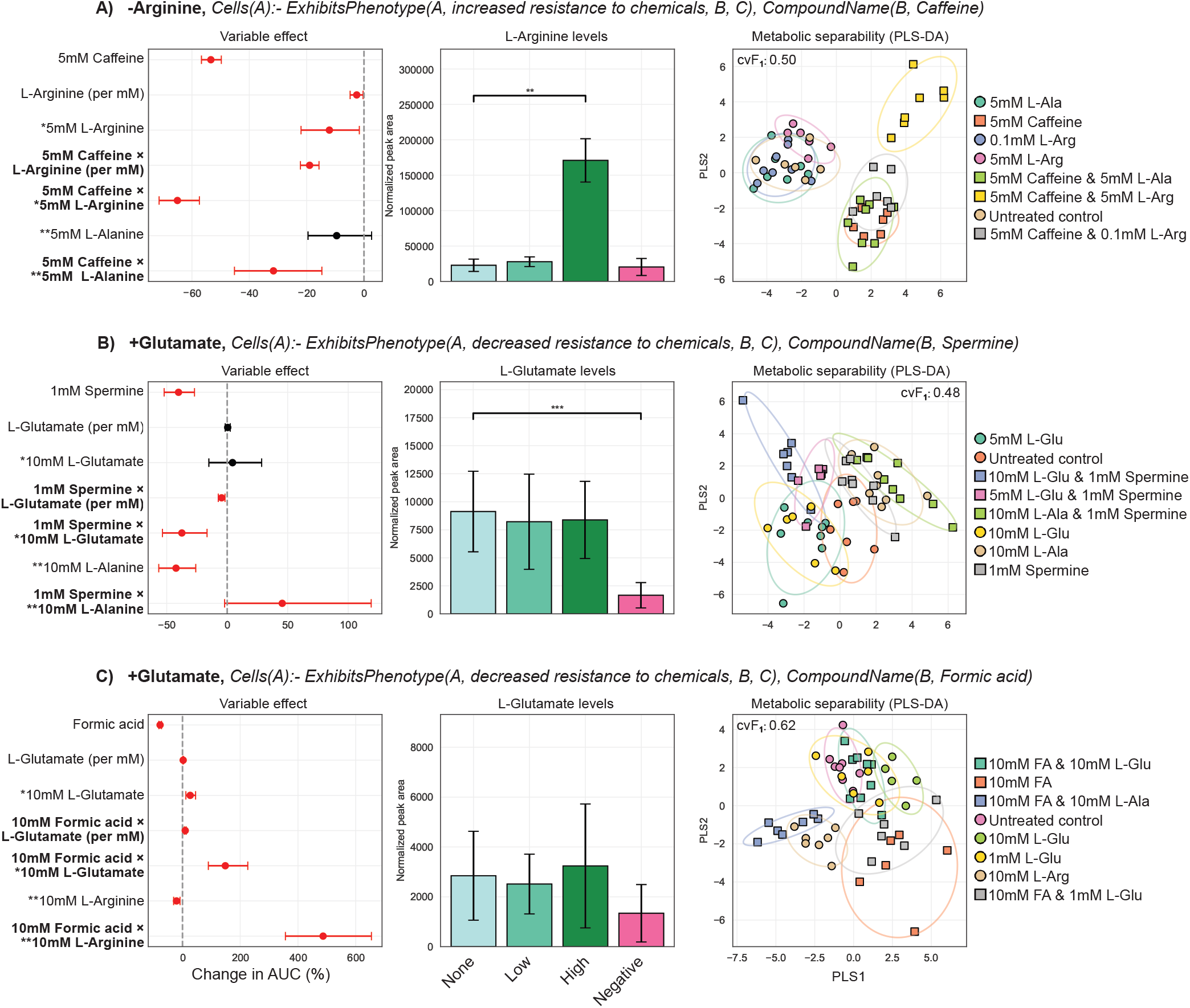
Results from automated interaction experiments with examples of different outcomes. The title of each row denotes logic program that was tested along with the learned association that defines the intervention. The forest plot signifies the coefficients and confidence intervals (2.5% and 97.5%) derived from a generalized linear model using AUC as the response variable. Red highlights significant (*p <* 0.05) variable effects as calculated by a non-parametric bootstrap (5000 resamples). The bar-plot shows the intracellular accumulation of the intervention compound, with the error-bars representing the standard deviation. *None, Low* and *High* denote the levels of concentrations of the intervention compound. The PLS-DA loading plot visualizes the two-dimensional separability of the metabolic profiles of the experimental groups, confirming a metabolic change from the supplied interventions. The classification performance for the full five PLS-DA components can be seen in the corner. *\*Effect scaled to correspond to an equimolar dose of the negative control. **Negative control*.

*Hypothesis 2* was partially consistent with empirical data (see Fig. 3B). Addition of only L-glutamate yielded a small—but statistically insignificant—increase in AUC. However, it significantly sensitized the cells to spermine (−4.6% per mM, *p <* 0.01). Intracellular L-glutamate accumulation did not increase with supplementation, suggesting a downstream metabolite may be the actual mediator of changes in resistance. The L-alanine control increased resistance to spermine (+45.5% at 10mM, *p <* 0.05) and significantly decreased L-glutamate accumulation (*p <* 0.001).

In a few select cases, the experimental validation of the hypothesis showed a non-predicted and non-specific interaction. For example, for *hypothesis 3*, as seen in Fig. 3C, the interaction effect between L-glutamate and formic acid was significant (+9.5% per mM, *p <* 0.001), but not in the predicted direction. The L-alanine negative control had a much larger effect on resistance under treatment compared to independent effects (+486% at 10 mM, *p <* 0.001), almost nullifying the effect of the stressor. L-glutamine and acetic acid (*hypothesis 4*) showed strong synergistic inhibition (−4.3% per mM, *p <* 0.05). However, L-leucine showed an equally strong effect (−36.9% at 10mM, *p <* 0.01), indicating the non-specificity of the intervention. Lglutamine was not significantly accumulated when supplemented, indicating a downstream mediator of reduced resistance.

Regarding *hypothesis 5*, L-lysine exhibited a specific, strong rescue effect against sucrose-induced hyper-osmotic stress (+8.9% per mM, *p <* 0.001), while valine produced a modest rescuing effect (+22.5% at 10 mM, *p <* 0.05). Intracellular L-lysine levels increased under the highest supplementation level (*p <* 0.05).

*Hypothesis 6* suggested a rescuing effect of L-arginine supplementation on lithium-induced stress, but this was inconsistent with the data. Exposure to 100 mM lithium chloride severely reduced AUC (−84.6%, *p <* 0.001). The intervention had little effect on resistance outcome, although it did increase intracellular L-arginine accumulation at higher levels of supplementation (*p <* 0.05). The L-alanine negative control showed a synergistic decrease in AUC relative to independent effects (−15.9% at 5mM, *p <* 0.05) and decreased intracellular concentrations of arginine (*p <* 0.05). *Hypothesis 7* (involving L-proline and lactic acid) showed minimal impact on treatment outcome. Both the intervention and the negative control affected growth dynamics moderately, but only the L-alanine negative control reached statistical significance (−38.6% at 20mM, *p <* 0.001).

All outcomes—whether supporting or refuting the initial hypothesis—are systematically recorded in a graph database.

Partial least squares discriminant analysis (PLS-DA) revealed clear separation of controls, interventions and stressors through their metabolic profiles. Cross-validation metrics indicated high model reliability, underscoring the consistency and specificity of the metabolomics data. Evaluation metrics and low dimensional representations of several investigations can be seen in Fig. 3. Processed data is stored in the graph database and raw data is stored in online repositories (see details in *data availability*).

### Hypothesis refinement showed alleviation of formic acid stress by aminoadipate

Regularized linear regression was applied to data from *hypothesis 3* (visualized in Fig. 3B) to model growth dynamics as a function of formic acid-treated metabolic profiles. By interpreting the model’s feature weights, this iterative analysis refined the initial hypothesis and led to a new, testable prediction. Based on the resulting ranked metabolites (see Fig. 4A), a logic program was constructed using experiment metadata and the highest-ranked locally available compound—aminoadipate (see below).

**Fig. 4.**
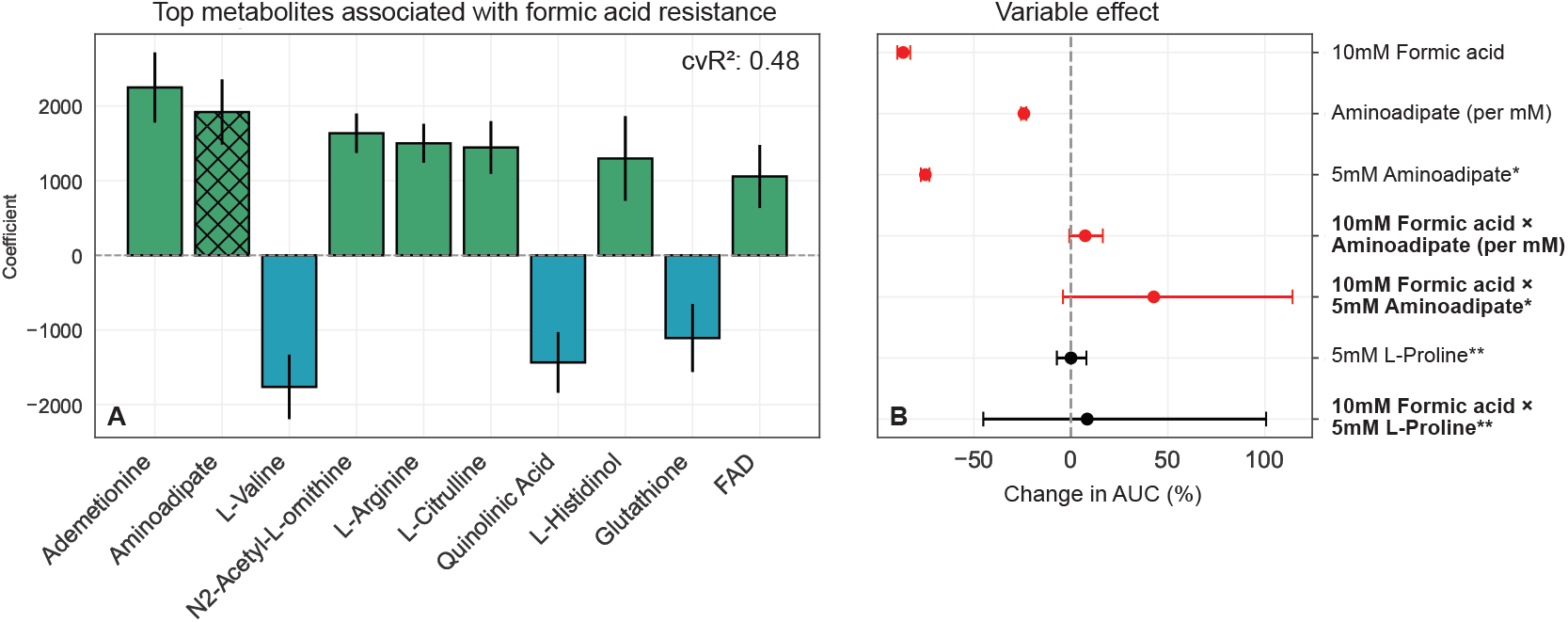
Iterative hypothesis refinement. **A**. The bar-plot denotes the average magnitude of coefficients derived from a linear regressor when predicting for resistance to formic acid stress (using data derived from *hypothesis 4*, see Fig. 3B). Regression performance is presented as coefficient of determination across repeated cross validation (*cvR*^2^). The hatched bar is the compound selected to generate the experiment (aminoadipate). The error bars denote the standard deviation across cross validation folds. **B**. The forest plot signifies the coefficients and confidence intervals (2.5% and 97.5%) derived from a generalized linear model using AUC as the response variable. Red highlights significant (*p <* 0.05) variable effects as calculated by a non-parametric bootstrap (5000 resamples). The interaction terms are of special importance to the hypothesised logic program. *\*Effect scaled to correspond to an equimolar dose of the specificity control. **Negative control*.

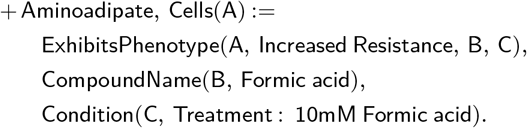

The hypothesis was fed into the initial step of the pipeline, and the experiment was subsequently performed. Results can be seen in Fig. 4B. Complete regression coefficients can be found in Sup. Table 3.

The results showed a partial rescue of formic acid stress (+7% per mM, *p <* 0.05) when combined with aminoadipate, confirming the hypothesis. Note that the supplement and treatment are independently highly toxic to the cells (see Fig. 4B), and as such, the dynamics might be subject to saturation effects.

### A database of hypotheses and experimental data enables the efficient reuse of experimental data

Although our automated procedures improve efficiency, conducting experiments remains costly. To enable reuse of empirical data for initial assessment of hypotheses, and to fully record the hypotheses and experimental data, existing ontologies (e.g. APO) were extended and connected. We then transformed the generated hypotheses, the experimental protocols, and the empirical data into OWL-DL statements and recorded these in a graph database (Fig. 5AB), which can be queried for empirical data relevant for a particular hypothesis. The following hypothesis—suggesting an interaction between intracellular L-proline accumulation and resistance to formic acid stress—was generated by the initial stages of the automated procedure (see *hypothesis 9* in table 1).

**Fig. 5.**
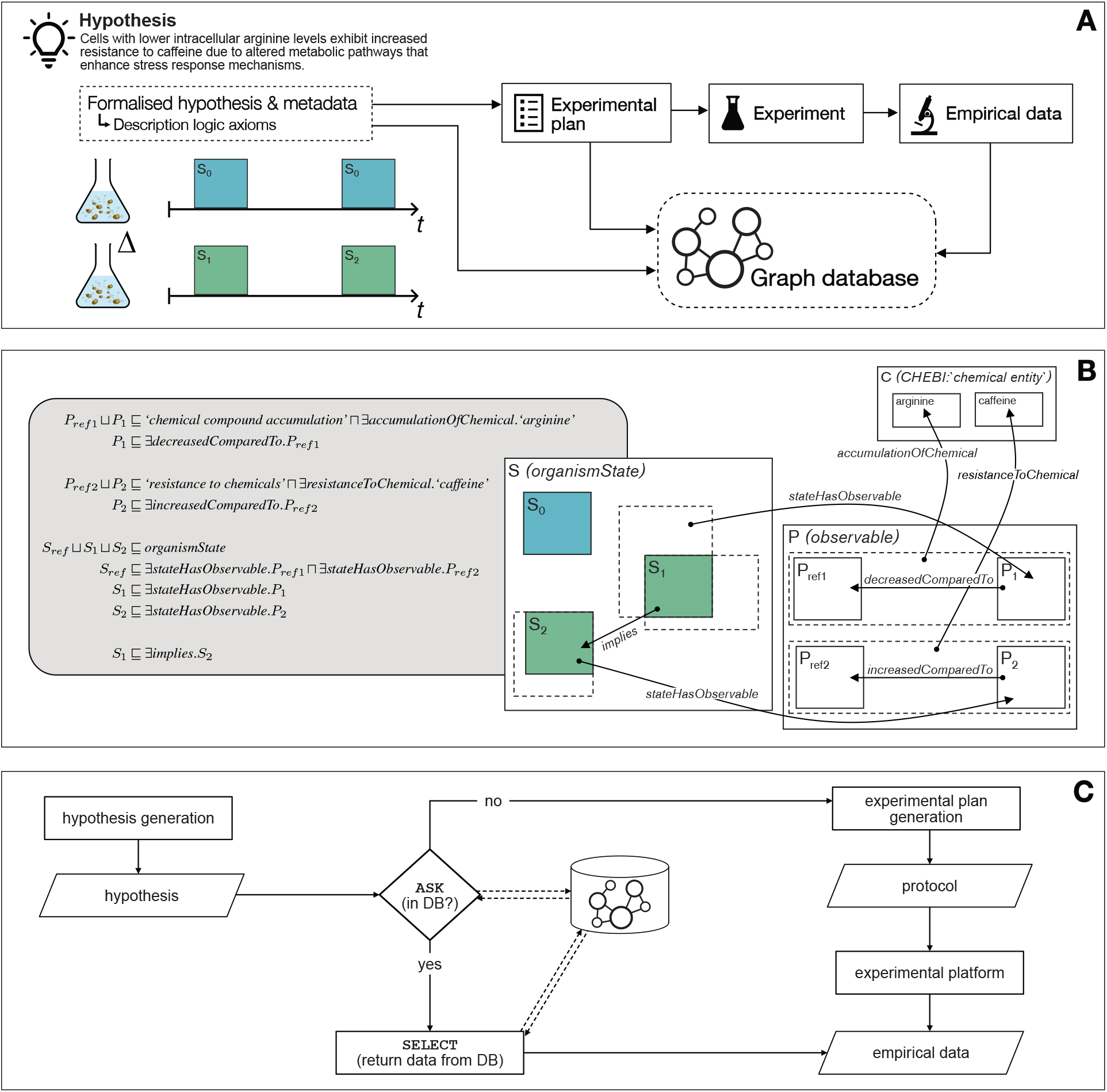
Graph Database and description logics. **A**. The hypothesis, in its original form as a logic program and also with the LLM output, is formalised into description logic axioms using an extended version of the Ascomycete Phenotype Ontology (APO); the generated experimental plans are stored using terms from the Ontology for Biomedical Investigations (OBI); and empirical data is stored using terms from the Genesis ontology. **B**. *(L)* DL axioms expressing the same hypothesis shown in A; *(R)* a visual representation of the concept inclusions expressed in the statements (with a subset of the relations shown, for clarity). **C**. To make efficient use of data already collected, an ASK SPARQL query is executed on the graph database to identify if there are empirical data that can be relevant to the hypothesis; if yes, then a SELECT query is executed to return the data, else if no, the full automated experimental generation procedure is followed.

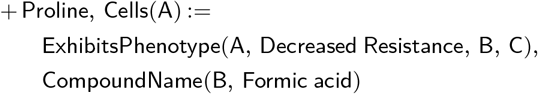

Before executing any experiments, the database was queried for relevant empirical data. Proline was used as a negative control for *hypothesis 8*, meaning there are data from experiments conducted with proline supplementation and formic acid treatment. Returning these data from the database— see Fig. 5C—shows no decrease in resistance to formic acid stress, see Fig. 4B, which is inconsistent with the hypothesis. Therefore, this hypothesis can be de-prioritised for experimental validation, conserving valuable resources.

## Discussion

The future of research lies in automated solutions. Advances in robotics, machine learning, and high-throughput technologies are transforming the scientific process (1, 3). These innovations help minimise human bias, accelerate research cycles, and reduce variation introduced by human error and incomplete recording of experimental protocols, environmental conditions, and other subtle factors that can influence outcomes (36–38). In this automated paradigm, algorithms design experiments, robots execute protocols, and the results are all automatically analysed, thus enabling exploration of vast experimental spaces beyond human throughput (4, 7, 8, 13). By codifying workflows and enforcing rigorous data-handling and reusability practices, such platforms ensure that every result is traceable, reproducible, and extensible. Ultimately, combining complex exploration with automated, quality-controlled systems promises to transform biology into a truly scalable, collaborative, and reliable discipline.

Building on this vision, we designed and evaluated an automated multi-agent framework that systematically produces and experimentally evaluates logic-derived, data-driven computational hypotheses, with metadata captured at every stage. We also demonstrate the utility of metabolomics data for evaluating interventions and for suggesting alternative explanations and follow-up experiments. Unlike typical agentic workflows—that orchestrate flexible yet fragile toolchains with limited semantic structure—our framework formalizes knowledge representation from the outset. This foundation enables consistent hypothesis generation, rigorous tracking of experimental logic, and clearer links between data and interpretation.

Several hypotheses generated by our framework point to biologically relevant but previously underexplored interactions in *S. cerevisiae*. One example is the interaction between L-glutamate and spermine, despite these being two essential compounds connected to each other via several different reactions in glutathione metabolism (KEGG) (39).

Another previously uncharacterized finding is the interaction between L-arginine and caffeine, likely mediated by distinct regulation of the TORC1-complex. Caffeine acts as an inhibitor of TORC1 in yeast, and arginine is tightly linked to TORC1 activity (40, 41).

Even when outcomes diverged from predictions, the data revealed significant interactions between interventions and stressors—for example, between L-lysine and sucrose-induced hyper-osmotic stress. Lysine related genes are downregulated under this stressor (42), but direct functional links remain sparse.

Through iterative hypothesis refinement, we demonstrate for the first time that aminoadipate confers resistance to formic acid stress, whereas L-glutamate’s role in modulating tolerance has been reported previously (43).

Testing these hypotheses highlighted mechanistic assumptions that proved limited. For example, treatments intended to increase intracellular glutamate sometimes failed to do so. In other cases (e.g., *hypotheses 2* and *3*), resistance phenotypes were likely driven by downstream metabolites rather than the supplemented compounds themselves. Interventions based on suggested metabolite concentrations may still modulate upstream or downstream reactions, explaining the disconnect between predicted concentrations and observed phenotypes (44). To capitalize on these insights, by archiving all outcomes under existing ontologies, the framework builds a knowledge base that supports future systems biology research and cross-study insights. Thus, the platform treats falsification as a valuable contribution rather than an uninformative failure.

The hypotheses generated and evaluated in this work are simple, but the framework can easily be extended to handle other types of hypotheses, such as those based on observed metabolite accumulations, changes in metabolite excretion rates, environmental stressors and drug-to-metabolite connections (relations present in the initial hypothesis generation steps, see Sup. Note 3). Its modular design also allows incorporation of additional data types, like transcriptomics and proteomics, with only minimal adjustments to the hypothesis-generation pipeline (45).

Although no intellectual input is needed from humans beyond defining the framework’s scope and limitations, the workflow is not fully automated: it still requires minimal human intervention for transferring sample plates between work stations. We also note that these actions are trivially automated should the method need to be scaled; moreover, adding a more comprehensive orchestration layer could coordinate task scheduling, making it truly autonomous.

Looking forward, scaling this automated workflow would require strict selection of hypotheses to maximize information gain per run. Integrating richer logic, decision-theoretic frameworks, econometric modelling and metrics from information theory could allow the system to better formulate and prioritize experiments, saving resources and contributing to scientific knowledge in a faster and more ethically sound manner (1, 8, 46). More advanced LLMs, such as complex reasoning models, could also allow more sophisticated hypotheses to be explored.

In this work, we demonstrate a flexible multi-agent framework for end-to-end hypothesis generation and experimental validation—combining LLMs, relational learning, and automated laboratory workflows. Our approach leverages mass-spectrometry metabolomics—scalable, low-cost and automation friendly—to provide a rich information source to drive discovery, and deliver high-throughput, reproducible insights, firmly rooted in logic and community-adopted ontologies. We believe this platform lays the groundwork for a more reliable, machine-driven discovery process in systems biology, operating at unprecedented scale and rigour.

## Methods

### Frequent pattern mining

The relational database used for hypothesis generation makes use of the Saccharomyces Genome Database (SGD), ChEBI, STITCH-DB, and the Yeast consensus model (Yeast9) as sources of relational data. All of the data used to create the database were downloaded using either AllianceGenome, STITCH-DB, or the Metabolic Atlas (all retrieved 2024-12-09) (27, 29–31, 47, 48).

The database consists of relations covering exhibited phenotypes of single gene deletant mutants and ontological descriptors of genetic and chemical perturbations. Additionally, it contains relations regarding protein-chemical interactions, and metabolism connected to the genes of interest (see Sup. Note 3 for a list of available relations). Only observables that are actionable within our experimental capacity were considered. This includes descriptors such as resistance to chemicals, compound accumulations and environmental sensitivities. Relations were filtered to only include facts derived from systematic mutation sets. The database is represented in Datalog, as its declarative structure enables clear and concise expression of biological relationships (34).

In order to extract biologically relevant patterns from the constructed database, relational learning was applied in the form of frequent pattern mining. These patterns were learned using the induce features mode in the ILP-engine aleph (v.5), utilizing a simplified version of the data-mining algorithm WARMR (25, 26). It is used to find frequent relational patterns using the deleted genes as positive examples, as in Brunnsåker et al. (45). The 25 genes with the most recorded phenotypes were used to construct the bottom-clause for the search (a bottom clause is the most specific clause that covers a given example(s), serving as a starting point for generalization). Complete parameters and a more detailed description of the search can be found in Sup. Note 3.

The resulting patterns were propositionalised into a tabular dataset of logic programs covering 4678 single-gene deletants from the Mülleder et al. metabolomics dataset (32). When multiple propositionalised programs described the same examples, we prioritized the most detailed ones using a scored vocabulary.

### Hypothesis weighting

In order to link logic programs with a biological observable, the propositionalised logic programs were used as independent variables for predicting amino acid accumulations, as measured in Mülleder et al. (32). Associations were learned using an ElasticNet algorithm (49). Note that this makes the implicit assumption that it is an accumulation that underlies the hypothesis, not the functional flux. Learned coefficients were then averaged across 10-fold cross-validation. Each logic program was tied to the coefficients, providing a weight (and a testable association) for each logic program and amino acid present in the dataset. Logic programs with zero-valued coefficients are removed from the pool of hypotheses for each metabolite observable. Coefficients were scaled to unit ∞norm. Amino acids with a negative *cvR*^2^ were removed from further consideration. Example hypotheses can be seen in the results.

### Semantic recording of hypotheses

The generated hypotheses have their origins in data downloaded from the Saccharomyces Genome Database (SGD), which records data using terms taken from ontologies: phenotypes are recorded using terminology defined in the Ascomycete Phenotype Ontology (APO) (27, 28), and chemical species are defined using terms from Chemical Entities of Biological Interest (ChEBI) (31).

APO was designed to record hypotheses about genotype– phenotype relations, and statements generally have four components: (1) the type of mutant; (2) the observable phenotype; (3) a qualifier on this observable (e.g. “increased”); and the experiment type^1^. The implicit definition of qualifiers in APO is for a mutant with respect to a wild-type strain, and is therefore insufficient for recording hypotheses concerning arbitrary changes in genotype or phenotype. We extend APO to include qualifying relations between collections of genotype and phenotype changes, called *organism states*. We also created relations to specify phenotypes that relate to chemical species, defined externally in ChEBI. These extensions are necessary to express the types of hypotheses handled in this work, for example: “cells with lower intracellular arginine levels exhibit increased resistance to caffeine.” Fig. 5A,B shows how this specific hypothesis is modelled using our framework.

The basic form of hypotheses in systems biology is *A* → *B*, and for a hypothesis to be testable, *A* should be an intervention achievable with the available laboratory equipment and consumables, and *B* is measurable. In our new terminology, *A* and *B* are *organism states*. The specific nature of the implication, *A → B*, will vary in different domains, and as such the *implies* relation should be seen as a top-level term. In this work, given the prompt provided to the LLM, we can interpret *implies*(*A*, *B*) as

> intervention to achieve state *A* will lead to an observation of state *B*, where an absence of intervention *A* will not[, and optionally where an alternative intervention *A*^*′*^ (negative control) also will not lead to an observation of state *B*.]

For now we delegate the assessment of the truth of this relation for each hypothesis to the statistical analyses described.

The axioms introduced to extend APO were edited using Protégé (v5.6.4) (50). We transform each hypothesis to OWL-DL statements using Python scripts. The resulting quads are stored in datasets on Apache Jena Fuseki server (v.4.5.0), building on the database system developed by Reder et al. (51). An example set of description logic axioms that record the hypothesis stated above, and the resulting OWL-DL statements (serialised in TriG format), is provided in Sup. Note 6.

### Generative experimental design

A standardized context (background, protocols and lab constraints) was used to prompt an LLM-agent for an initial, detailed explanation of a generated hypothesis and a basic experimental plan to test it. A negative control is automatically selected from a list of amino acids that have zero-valued coefficients for the generated hypothesis. The contexts, explanation and prompts are saved in the graph database for each hypothesis. Templates can be found in the GitHub repository (see *code availability*).

To ensure increased robustness, several hypothesis-plan variations are then generated with the same context, and a second LLM-agent picks the best one based on relevance to yeast physiology, safety and context fidelity. Note, that in order to comply with safety regulations of the laboratory setup in use, this final experimental plan is at this stage manually assessed for safety before being passed on.

A third LLM-agent auto-formalises the chosen experimental plan into a JSON-file, which is then validated against the original context and hypothesis. If it fails any check, the JSON is regenerated up to three times. Experimental plans are generated and auto-formalised via the OpenAI API using the GPT-4o model (v.2024-08-06) through the official Python client library (openai v.1.44.1) (33).

Using the formalised, machine-readable experimental protocol, the experimental variables are automatically identified, and a robust plate-layout is generated using PLAID (52). Using the layout, a library of stock compounds and the experimental protocol, a dispensing scheme with aspiration and dispensing targets and volumes for use with Hamilton Venus Software (v.5.0.1; Hamilton Company) is generated. An example scheme can be seen in Sup. Table 4.

A mass spectrometry run-list is automatically produced from the supplied protocol and user-defined parameters. The injection order is randomized to mitigate instrumental variation and systematic effects. Study quality control samples (sQCs) were placed systematically across the entire run-list, before and after biological samples, allowing for normalization and batch correction. Blocks of samples were separated by blank (extraction solvent) injections in order to assess analyte carry-over and background signal. An example run-list can be seen in Sup. Table 5.

### Automated sample preparation and cultivation

The generated plate-layout and dispensing schemes are executed on a Hamilton Microlab Star (using pre-prepared stocks of the available compounds) producing the microwell plate used for the experiment. Unless otherwise specified, experiments use minimal YNB medium (with ammonium sulfate, 2% glucose, and 75 µg/mL ampicillin, but no amino acids). The strain used to represent a reference genotype was selected to be Δ*ho* from the prototrophic strain collection constructed in Mülleder et al. (53)—due to the inertness of the deletion in non-mating-type contexts.

Each of the generated hypotheses were tested on separate microwell plates, allowing for up to 10 biological replicates per experimental condition, depending on the suggested intervention. Intervention and perturbation compounds present in the library are dissolved in sterile milliQ-water.

To run the cultivation and sampling, an Overlord (v.2.0) script (the orchestration software used to control the robotics and instrumentation in Eve) is generated programmatically using predefined parameters, such as cultivation time and plate-reader configurations (4, 5, 8). For all of the experiments performed in this project, cultivation duration was set to 20h. See Sup. Note 2 for a complete description of the cultivation protocols used.

### Automated sample processing

Sampling for mass spectrometry analysis is automatically performed after experiment termination. Quenched samples are prepared by robotically passing the cultivation plate to a Bravo Automated Liquid Handling Platform and transferring the samples into −80°C methanol (99% purity). The ratio between sample and methanol is kept at 1:1 v/v. The quenching plate is then subsequently passed to a centrifuge, where it is centrifuged at 2240g for 10 minutes. The supernatant is then discarded and the pellet used for subsequent metabolite extraction.

Metabolite extraction is performed using a standardized, automated variation of the extraction protocol used in Brunnsåker et al. (5). Extraction is robotically performed in a Hamilton Microlab Star, where 150uL of 80% ethanol (preheated to boiling temperatures) is dispensed into the sample plate, which is placed on a Hamilton HeaterShaker add-on set at 100°C to avoid drops in temperature. The sample plate is shaken vigorously at 1600 rpm for 2 minutes to expedite resuspension of the quenched cells. The supernatant is transferred to a filter plate (0.2uM, VWR, 97052-124) and separated using positive pressure filtering (set at 40psi for 60 seconds).

Study QCs are prepared manually through overnight cultivation of Δ*ho* in reference conditions (30°C, YNB without amino acids). Samples are quenched in 1/1 (v/v) −80°C pure methanol, and subsequently, extracted in 80% ethanol heated to boiling temperatures through a water bath. The boiling ethanol is poured over the cells, and vortexed for 1 minute, and put back in the hot water bath for 4 minutes. The pellet is discarded, and the supernatant is stored in −80°C pending analysis. This is similar to protocols used in Brunnsåker et al. and Reder et al. (22).

### Growth metrics and hypothesis testing

Growth curves are constructed from raw optical density (OD) readings at 600nm using a BMG Omega Polarstar. The growth curves are then automatically processed to remove outliers. Outlier detection is based on the median absolute deviation (MAD) of log-transformed area under the curve (AUC), total variation and final OD values. Wells exceeding a threshold of 3 MADs for any parameter are identified, excluded and reported. Curves are normalized to background by subtracting the average value of the three closest blanks on the microwell plate. The remaining curves are smoothed using locally estimated scatterplot smoothing (LOESS, statsmodels, v.0.14.4) (54). AUC is calculated using the composite trapezoidal rule on the smoothed curves (numpy, v.1.24.3) (55).

The impact of treatment and intervention on growth dynamics are automatically tested using log-transformed AUC as the response variable, and fitting a generalized linear model (statsmodels, v.0.14.4) that included terms for treatment, the concentration of the intervention, an interaction term, and a separate flag for negative controls. To account for skewness and unequal variances, a non-parametric bootstrap (5000 resamples) is used to compute 95% confidence intervals and empirical p-values for each main and interacting effect. Example results can be seen in Fig. 3. The experimental variables, such as concentrations, are extracted from the auto-formalized protocols.

### Automated metabolomics data acquisition and analysis

An internal Collision Cross-Section (CCS) library was created using metabolites present in the Yeast9 Genome scale metabolic model as compounds of interest (29). CCS-values for relevant metabolites were acquired from AllCCS (56). In case of missing experimental records for common adducts of amino acids (as these are the main targets of the generated hypotheses), predicted CCS values from AllCCS were used.

Mass spectrometry analysis is performed using an Agilent RapidFire-365 and an Agilent 6560 DTIMS-QToF system (23, 24). The instrument Data acquisition and peak picking is done automatically via AutonoMS, using a run-list generated from the hypothesis metadata and experimental protocol as the input (22). Peaks are identified based on both m/z (mass-to-charge ratio) and CCS (*Å*^2^) (57). The cartridge in use for all experiments was HILIC (Hydrophilic Interaction Liquid Chromatography). Details regarding mobile phases, method parameters, acquisition settings, demultiplexing parameters and peak picking parameters can be seen in Sup. Note 4 and Sup. Note 5.

The AutonoMS-output (a tabular representation of identified metabolites with an associated peak area) is initially curated using sensor outputs from the RapidFire system, removing injections with insufficient injected sample (≥600*ms*). Peaks with an average area below the included blanks (extraction solvent), along with peaks with more than 1/3 missing detections across study samples, are excluded from further analysis. In case of several identified adducts per metabolite, the one with the lowest relative standard deviation (RSD) in the sQC samples is used. Samples are normalized using Probabilistic Quotient Normalization (PQN), utilising the sQC-samples as the reference spectra (58). Missing values are imputed with multivariate iterative imputation using RandomForest (59). Samples outside of the 95% Hotelling T2 ellipse (using PCA-derived principal components) and high unexplained residual variances (beyond 99th percentile) are treated as outliers, similar to quality control recommendations suggested by González-Domínguez *et al*. (60). Outlier reports are included in the experiment summaries found in the GitHub repository.

To estimate the validity of the LLM-suggested interventions (confirming intracellular accumulation), pairwise comparisons of the target metabolites across untreated conditions is performed using a Wilcoxon rank-sum test (scipy, v.1.15.1) (61). Metabolic separability is classified using (5-component) partial least-squares discriminant analysis (PLS-DA), treating each experimental group as a separate class (mixOmics, v.6.22.0) (62). Classification performance is evaluated using the macro *F*_1_ score.

Metabolites associated with resistance to treatment are determined with ElasticNetCV, using repeated cross-validation (scikit-learn v.1.2.2) (63). AUC of treated samples are used as the dependent variable. Example results can be seen in Fig. 3.

As part of the evaluation, metabolomics data were collected for each individual cultivation in the study, producing 976 separate biological injections, including 336 sQC injections. The metabolomics dataset spans 52 unique conditions, with 21 different nutrient compositions, and 8 treatments, including controls. Each condition has a minimum of 10 biological replicates and associated growth dynamics. See *data availability* for details.

### Linking hypotheses and experimental plans to data

We also record the experimental plans, and the resulting empirical data from completed experiments, using controlled vocabularies in a graph database. These plans and data are linked to the hypothesis through a combination of terms from existing ontologies and new terms. In the case of experimental protocols the terms are taken primarily from the Ontology for Biomedical Investigations (OBI) (64); and terms for growth and metabolomics are taken primarily from the Genesis ontology (51).

The JSON files containing the experimental protocols, and the tabular files containing growth and metabolomics data, are then mapped to relevant terms from existing ontologies using RMLMapper (v7.3.3); the mapping files can be found in the GitHub repository (see *code availability*).

## Data availability

All data are included in the supplementary information or available from the authors. Mass-spectrometry data and associated metadata can be downloaded from Zenodo at https://doi.org/10.5281/zenodo.15730908.

## Code availability

Code (including cultivation, quenching and extraction scripts), contexts, complete supplemental information and experimental results can be found on GitHub at https://github.com/DanielBrunnsaker/GenExp.

## ACKNOWLEDGEMENTS

The authors gratefully acknowledge the members of the Ross King Group at Chalmers University and Cambridge University for their insights and discussions. We also thank the members of the Chalmers Mass Spectrometry Infrastructure for their help and technical expertise. Additionally, we thank the Ralser Lab at Charité–Universitätsmedizin Berlin for providing the yeast strains used in this study. This work was partially supported by the Wallenberg AI, Autonomous Systems and Software Program (WASP) funded by the Knut and Alice Wallenberg Foundation

## AUTHOR CONTRIBUTIONS

D.B. and R.D.K. conceptualized the study. D.B. and A.H.G. designed the generation and analysis framework. D.B. and P.N. designed the experimental pipelines and the integration between platforms. A.H.G., F.K. and D.B. designed the ontologies and databases. D.B., P.N. and E.Y.B. executed the experiments. D.B., E.Y.B. and I.A.T. analysed the results. D.B., A.H.G. and R.D.K. wrote the manuscript. All authors reviewed the manuscript.

^1^Note that experiment type will be absent in the case of a hypothesis not yet linked to any empirical data.

