## Supplemental Notes for "Agentic AI Integrated with Scientific Knowledge: Laboratory Validation in Systems Biology"

### Self-driven Biological Discovery through Automated Hypothesis Generation and Experimental Validation

July 20, 2025

### Supplementary Note 1: Overview of hypothesis space

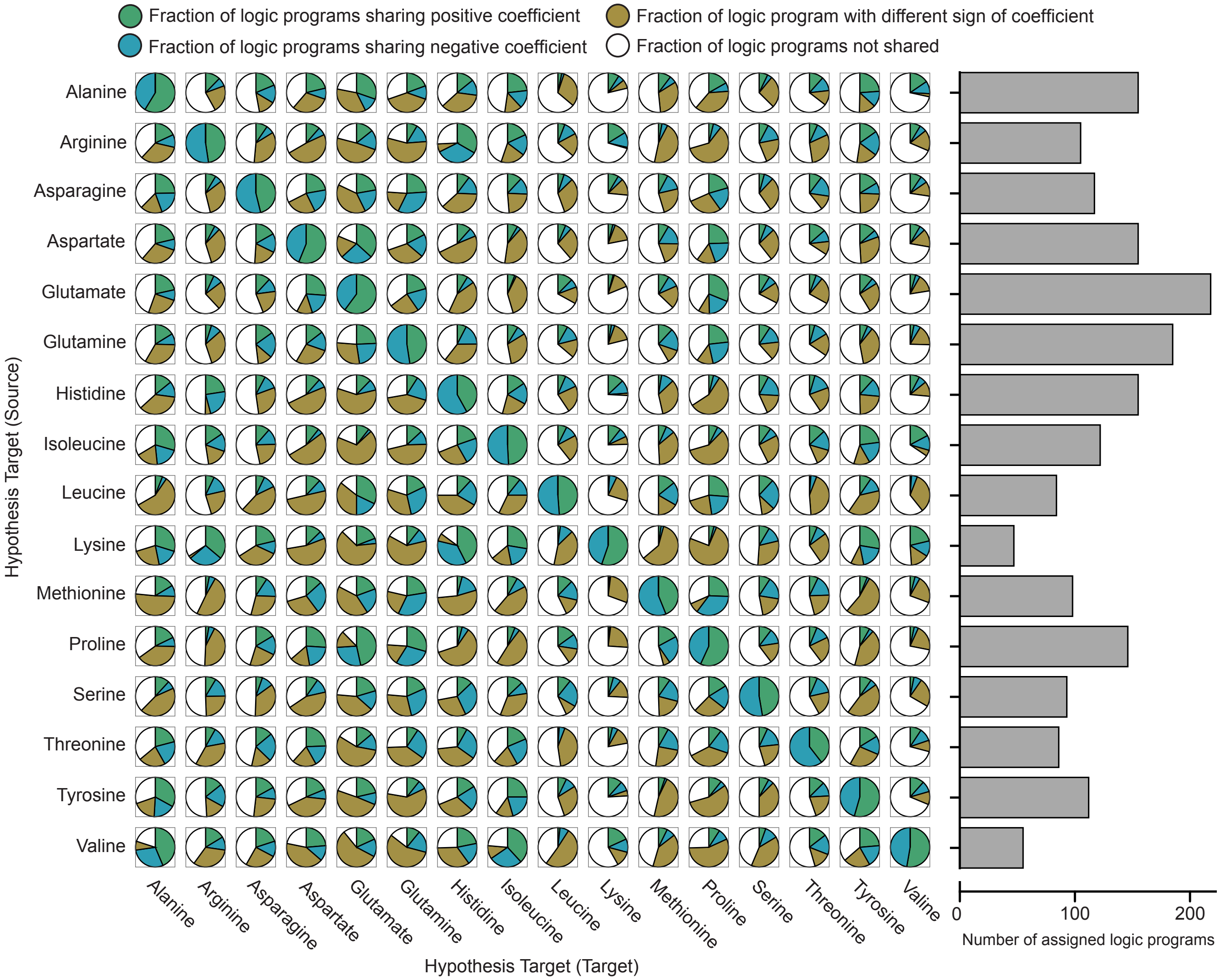


**Fig. 6.** Asymmetric hypothesis-space overlap between pairs of amino acids. Each pie chart shows—as a fraction of the total number of logic programs that are assigned to the amino acid on the vertical axis—the fraction of logic programs that also are assigned to the amino acid on the horizontal axis. For those that are shared, the blue, green, and yellow segments represent respectively positive, negative, and differing signs of coefficient in the linear regression model for prediction of abundance. The bar chart shows the total number of assigned logic programs for each amino acid on the vertical axis.

### Supplementary Note 2: Cultivation procedures

After plate preparation in a Hamilton Microlab Star, growth profiling was performed using the automated laboratory cell "Eve". The plate is transferred from Eve's Cytomat™ automated incubator (30°C) to a Teleshaker Magnetic Shaking System, where it is shaked for 30s at 800rpm, divided evenly between clockwise and counter-clockwise double-orbital shaking. After shaking the plate is transferred to a BMG Polarstar plate reader, where it undergoes optical density measurements at 600nM (the temperature in the plate reader was kept at a constant 30°C). After measuring, the plate is returned to the incubator. The protocol is automatically repeated every 20min for 20h (or the time specified in the generated protocol). At experiment termination, the microwell plate is transferred to an Agilent Bravo liquid handling station, depending on user preference.

Runtime-logs were saved for all of the cultivations done in the study. Note that in some cases several plates were run in parallel, slightly complicating the interpretation of the logs. These records—and others—can be found in the accompanying GitHub repository or by querying the graph database.

### Supplementary Note 3: Allowed relations & Search parameters

In order to extract valid logic programs from the constructed database, relational learning was applied in the form of frequent pattern mining using a simplified version of the data-mining algorithm WARMR in aleph (version 5, https://www.cs.ox.ac.uk/activities/programinduction/Aleph/aleph.html). The search was performed using the deletant strains provided in (23) as positive examples. WARMR is a data mining algorithm that uses a level-wise search algorithm, iteratively adding logical conditions until some constraint (defined by the search parameters) has been achieved. When utisiling WARMR the search is restricted to a subset of first-order logic (due to the difficulties of propositionalising higher-order patterns).

The relations allowed in the parameter search are the following:

$$\begin{aligned}
&\mathsf{Genotype}(+\mathsf{ORF})\\
&\mathsf{ExhibitsPhenotype}(+\mathsf{ORF},\ \#\mathsf{State},\ -\mathsf{Value},\ -\mathsf{Cond}),\\
&\mathsf{CompoundName}(+\mathsf{Value},\ \#\mathsf{Name}),\\
&\mathsf{CompoundModulatesTarget}(+\mathsf{Value},\ \#\mathsf{Action},\ -\mathsf{ORF}),\\
&\mathsf{ParticipatesInMetabolism}(+\mathsf{ORF},\ \#\mathsf{Type},\ \#\mathsf{Metabolite}),\\
&\mathsf{Condition}(+\mathsf{Cond},\ \#\mathsf{Description})
\end{aligned}$$

The parameters used for the pattern-search in Aleph, can be seen below, along with a short description.

| **Parameter** | **Value** | **Description** |
|---|---|---|
| *i* | 20 | Upper bound on layers of new variables |
| *clauselength* | 4 | Upper bound on number of literals in an acceptable clause |
| *search* | ar | Uses WARMR as the basis of the search |
| *pos_fraction* | 0.001 | Rules must cover at least this fraction of the total examples |
| *max_features* | 4097 | Upper bound on the maximum number of boolean features |
| *noise* | inf | Upper bound on the number of negative examples allowed |
| *nodes* | 200000 | Upper bound on the nodes to be explored when searching for an acceptable clause |

### Supplementary Note 4: Parameters for metabolomics acquisition

The RapidFire method used typically involved a sample sipping time of 600ms, followed by a loading-phase lasting 4000ms. It was then followed by a 7000ms sample elution into the MS, and lastly, 1500ms for equilibration. The sipper was washed between injections in organic (Methanol) and acqeous (Water) solvents. One injection has an analysis-time of 13.1 seconds.

Mobile phases were prepared with 10 mM ammonium formate and 0.176% formic acid. Phase A comprised 90% acetonitrile, 5% methanol and 5% water, while Phase B comprised 50% acetonitrile, 5% methanol and 45% water. The protocol in use was a modified version of the one derived from Mülleder et al. (23). Mass spectrometry data was collected in IM-QTOF mode with 4-bit multiplexed introduction of the ion packets into the drift tube. The Agilent 6560 was operated in the 100–1700 m/z range at a frame rate of 1.1 frames/s and a gas temperature of 325 °C. Analysed adducts include $[M+H]^+$, $[M+Na]^+$, $[M-H_2O+H]^+$ and $[M+NH_4]^+$. The mass spectrometer was run in positive mode.

**Table 2.** The Agilent 6560-IM-MS acquisition parameters used for the data collection in positive mode.

| Parameter | Value |
|---|---|
| *Ion Source* | Agilent Dual AJS ESI |
| *MS Abs. Threshold* | 200 |
| *Ion Mobility Mode* | IMS QTOF |
| *Component Model* | G6560B |
| *MS Rel. Threshold (%)* | 0.01 |
| *Min Range (m/z)* | 50 |
| *Max Range (m/z)* | 1700 |
| *Frame Rate (frames/sec)* | 1.1 |
| *IM Transient Rate (transients/frame)* | 14 |
| *Max Drift Time (ms)* | 60 |
| *Trap Fill Time (μs)* | 3900 |
| *Trap Release Time (μs)* | 100 |
| *Gas Temperature (C)* | 325 |
| *Gas Flow (l/min)* | 11 |
| *Nebulizer (psi)* | 45 |
| *Sheath Gas Temp (C)* | 275 |
| *Sheath Gas Flow (l/min)* | 12 |
| *VCap (V)* | 4000 |
| *Nozzle Voltage (V)* | 1000 |
| *Drift Tube Entrance Voltage (V)* | 1274 |
| *Drift Tube Exit Voltage (V)* | 224 |

### Supplementary Note 5: AutonoMS parameters

**Table 3.** AutonoMS method parameters. Encompasses both RapidFire method settings, preprocessing parameters and peak picking parameters.

| Parameter | Value |
|---|---|
| *Sipper Height (mm)* | 1 |
| *Wash Between Sips* | 1 |
| *No. Flushes After Plates* | 0 |
| *Pump1FlowRate (ml/min)* | 1.25 |
| *Pump2FlowRate (ml/min)* | 0.01 |
| *Pump3FlowRate (ml/min)* | 1.25 |
| *Pump 1 [B, C, D] Line %* | [100, 0, 0] |
| *Pump 2 [B, C, D] Line %* | [100, 0, 0] |
| *Pump 3 [B, C, D] Line %* | [100, 0, 0] |
| *Plate Configuration* | 96-well-plate |
| *Missed Sip Tolerance* | 10000 |
| *Aspirate Cycle Duration (ms)* | 600 |
| *Load/Wash Cycle Duration (ms)* | 4000 |
| *Extra Wash Cycle Duration (ms)* | 0 |
| *Elute Cycle Duration (ms)* | 7000 |
| *Reequilibrate Cycle Duration (ms)* | 1500 |
| *Chromatography/infusion moving average* | 3 |
| *Minimum pulse coverage (%)* | 50 |
| *Moving average window smoothing size (drift)* | 3 |
| *Signal intensity lower threshold* | 5 |
| *Resolving power (IM)* | 30 |
| *Resolving power (TOF)* | 30000 |
| *High selectivity extraction* | Yes |
| *Method match tolerance (m/z)* | 0.005 |

### Supplementary Note 6: Example hypothesis in description logic

This is the arginine caffeine hypothesis, *hypothesis 1*.

#### A. DL Axioms.

$$P_{ref1} \sqcup P_1 \sqsubseteq \textit{'chemical compound accumulation'} \sqcap \exists \textit{accumulationOfChemical}.\textit{'arginine'} \quad \textbf{(1)}$$

$$P_1 \sqsubseteq \exists \textit{decreasedComparedTo}.P_{ref1} \quad \textbf{(2)}$$

$$P_{ref2} \sqcup P_2 \sqsubseteq \textit{'resistance to chemicals'} \sqcap \exists \textit{resistanceToChemical}.\textit{'caffeine'} \quad \textbf{(3)}$$

$$P_2 \sqsubseteq \exists \textit{increasedComparedTo}.P_{ref2} \quad \textbf{(4)}$$

$$S_{ref} \sqcup S_1 \sqcup S_2 \sqsubseteq \textit{organismState} \quad \textbf{(5)}$$

$$S_{ref} \sqsubseteq \exists \textit{stateHasObservable}.P_{ref1} \sqcap \exists \textit{stateHasObservable}.P_{ref2} \quad \textbf{(6)}$$

$$S_1 \sqsubseteq \exists \textit{stateHasObservable}.P_1 \quad \textbf{(7)}$$

$$S_2 \sqsubseteq \exists \textit{stateHasObservable}.P_2 \quad \textbf{(8)}$$

$$S_1 \sqsubseteq \exists \textit{implies}.S_2 \quad \textbf{(9)}$$

#### B. RDF/OWL Statements (TriG format).

```
@prefix hypo:  <http://hypo.project-genesis.io#> .
@prefix obo:   <http://purl.obolibrary.org/obo/> .
@prefix owl:   <http://www.w3.org/2002/07/owl#> .
@prefix rdfs:  <http://www.w3.org/2000/01/rdf-schema#> .
@prefix dct: <http://purl.org/dc/terms/> .

<http://hypo.project-genesis.io/hypothesis-metadata> {
    <http://hypo.project-genesis.io/H-0196d121-3310-771d-ab08-bb5f9f4973d4>
            dct:created  "2025-03-13T15:39:00"^^<http://www.w3.org/2001/XMLSchema#dateTime> ;
            dct:creator  "Genesis" .
}

<http://hypo.project-genesis.io/H-0196d121-3310-771d-ab08-bb5f9f4973d4> {
    hypo:S-0196d121-3400-715f-95a9-0430c564ffdb
            rdfs:subClassOf  [ a                   owl:Restriction ;
                               owl:onProperty      hypo:implies ;
                               owl:someValuesFrom  hypo:S-0196d121-3500-7ad8-bf51-ec89040bdb84
                             ] .
}

<http://hypo.project-genesis.io/states> {
    hypo:S-REF-0196d120-c332-7b22-b220-572b040a96a8
            rdfs:subClassOf  [ a                   owl:Restriction ;
                               owl:onProperty      hypo:stateHasObservable ;
                               owl:someValuesFrom  hypo:P-REF-0196d121-33e2-76a8-b698-330c8911c9ae
                             ] ;
            rdfs:subClassOf  [ a                   owl:Restriction ;
                               owl:onProperty      hypo:stateHasObservable ;
                               owl:someValuesFrom  hypo:P-REF-0196d121-34ed-799c-b5ea-296d85f551ce
                             ] .

    hypo:S-0196d121-3500-7ad8-bf51-ec89040bdb84
            rdfs:subClassOf  hypo:organismState ;
            rdfs:subClassOf  [ a                   owl:Restriction ;
                               owl:onProperty      hypo:stateHasObservable ;
                               owl:someValuesFrom  hypo:P-0196d121-34f7-7dc8-a499-7a7664f3e710
                             ] .

    hypo:S-0196d121-3400-715f-95a9-0430c564ffdb
            rdfs:subClassOf  hypo:organismState ;
            rdfs:subClassOf  [ a                   owl:Restriction ;
                               owl:onProperty      hypo:stateHasObservable ;
                               owl:someValuesFrom  hypo:P-0196d121-33f6-7e22-b286-acc2a7a6227e
                             ] .
}

<http://hypo.project-genesis.io/phenotypes> {
    hypo:P-REF-0196d121-33e2-76a8-b698-330c8911c9ae
            rdfs:subClassOf  obo:APO_0000095 ;                                # 'chemical compound accumulation'
            rdfs:subClassOf  [ a                   owl:Restriction ;
```

```
                                  owl:onProperty      hypo:accumulationOfChemical ;
                                  owl:someValuesFrom  obo:CHEBI_29016                # 'arginine'
                                ] .

    hypo:P-REF-0196d121-34ed-799c-b5ea-296d85f551ce
            rdfs:subClassOf  obo:APO_0000087 ;                                       # 'resistance to chemicals'
            rdfs:subClassOf  [ a                  owl:Restriction ;
                                  owl:onProperty      hypo:resistanceToChemical ;
                                  owl:someValuesFrom  obo:CHEBI_27732                # 'caffeine'
                                ] .

    hypo:P-0196d121-33f6-7e22-b286-acc2a7a6227e
            rdfs:subClassOf  obo:APO_0000095 ;                                       # 'chemical compound accumulation'
            rdfs:subClassOf  [ a                  owl:Restriction ;
                                  owl:onProperty      hypo:accumulationOfChemical ;
                                  owl:someValuesFrom  obo:CHEBI_29016                # 'arginine'
                                ] ;
            rdfs:subClassOf  [ a                  owl:Restriction ;
                                  owl:onProperty      hypo:decreasedComparedTo ;
                                  owl:someValuesFrom  hypo:P-REF-0196d121-33e2-76a8-b698-330c8911c9ae
                                ] .

    hypo:P-0196d121-34f7-7dc8-a499-7a7664f3e710
            rdfs:subClassOf  obo:APO_0000087 ;                                       # 'resistance to chemicals'
            rdfs:subClassOf  [ a                  owl:Restriction ;
                                  owl:onProperty      hypo:resistanceToChemical ;
                                  owl:someValuesFrom  obo:CHEBI_27732                # 'caffeine'
                                ] ;
            rdfs:subClassOf  [ a                  owl:Restriction ;
                                  owl:onProperty      hypo:increasedComparedTo ;
                                  owl:someValuesFrom  hypo:P-REF-0196d121-34ed-799c-b5ea-296d85f551ce
                                ] .
}
```

### Supplementary Note 7: Growth statistics from performed investigations

| Term | Estimate | CI 2.5% | CI 97.5% | % Change | p-value |
|---|---|---|---|---|---|
| Intercept | 11.0678 | 11.0075 | 11.1247 | | 0.0002 |
| Caffeine | -0.7636 | -0.8366 | -0.6883 | -53.4013 | 0.0002 |
| L-Arginine (per mM) | -0.02573 | -0.04959 | -0.003280 | -2.5404 | 0.01180 |
| L-Arginine at 5.0 mM | -0.1287 | -0.2479 | -0.01640 | -12.07287 | 0.01180 |
| Caffeine:L-Arginine (per mM) | -0.2103 | -0.2507 | -0.1707 | -18.9652 | 0.0002 |
| Caffeine:L-Arginine (at 5.0 mM) | -1.05146 | -1.2535 | -0.8533 | -65.05719 | 0.0002 |
| L-Alanine (at 5.0 mM) | -0.1000 | -0.2171 | 0.02621 | -9.5239 | 0.05179 |
| Caffeine:L-Alanine (at 5.0 mM) | -0.3794 | -0.6010 | -0.1589 | -31.5732 | 0.0010 |

**Table 4.** Summary statistics of *Hypothesis 1* (L-Arginine, caffeine and L-Alanine) with AUC of the time-series growth curve as the dependent variable in different treatment conditions.

| Term | Estimate | CI 2.5% | CI 97.5% | % Change | p-value |
|---|---|---|---|---|---|
| Intercept | 9.2949 | 9.1364 | 9.4773 | | 0.0002 |
| Spermine | -0.5147 | -0.7362 | -0.3152 | -40.2329 | 0.0002 |
| L-Glutamate (per mM) | 0.0043 | -0.0164 | 0.0251 | 0.4327 | 0.3413 |
| L-Glutamate (at 10.0 mM) | 0.0432 | -0.1643 | 0.2507 | 4.4117 | 0.3413 |
| Spermine:L-Glutamate (per mM) | -0.0469 | -0.0766 | -0.0182 | -4.5859 | 0.0012 |
| Spermine:L-Glutamate (at 10.0 mM) | -0.4694 | -0.7658 | -0.1822 | -37.4649 | 0.0012 |
| L-Alanine (at 10.0 mM) | -0.5499 | -0.8338 | -0.3010 | -42.2992 | 0.0004 |
| Spermine:L-Alanine (at 10.0 mM) | 0.3750 | -0.0216 | 0.7841 | 45.5027 | 0.0322 |

**Table 5.** Summary statistics of *Hypothesis 2* (L-Glutamate, spermine and L-Alanine) with AUC of the time-series growth curve as the dependent variable in different treatment conditions.

| Term | Estimate | CI 2.5% | CI 97.5% | % Change | p-value |
|---|---|---|---|---|---|
| Intercept | 10.7106 | 10.6006 | 10.8087 | | 0.0002 |
| Formic acid | -1.4950 | -1.6995 | -1.2983 | -77.5751 | 0.0002 |
| L-Glutamate (per mM) | 0.0236 | 0.0110 | 0.0374 | 2.3924 | 0.0010 |
| L-Glutamate (at 10.0 mM) | 0.2364 | 0.1105 | 0.3739 | 26.6715 | 0.0010 |
| Formic acid:L-Glutamate (per mM) | 0.0910 | 0.0640 | 0.1182 | 9.5214 | 0.0002 |
| Formic Acid:L-Glutamate (at 10.0 mM) | 0.9095 | 0.6398 | 1.1822 | 148.3068 | 0.0002 |
| L-Arginine (at 10.0 mM) | -0.2302 | -0.3673 | -0.0894 | -20.5651 | 0.0008 |
| Formic acid:L-Arginine (at 10.0 mM) | 1.7698 | 1.5185 | 2.0208 | 486.9951 | 0.0002 |

**Table 6.** Summary statistics of *Hypothesis 3* (L-Glutamate, formic acid and L-Arginine) with AUC of the time-series growth curve as the dependent variable in different treatment conditions.

| Term | Estimate | CI 2.5% | CI 97.5% | % Change | p-value |
|---|---|---|---|---|---|
| Intercept | 10.8973 | 10.8142 | 11.0033 | | 0.0002 |
| 30% sucrose | -1.9001 | -2.0451 | -1.7550 | -85.0445 | 0.0002 |
| L-Lysine (per mM) | -0.0828 | -0.1024 | -0.0637 | -7.9499 | 0.0004 |
| L-Lysine (at 10.0 mM) | -0.8284 | -1.0240 | -0.6370 | -56.3241 | 0.0004 |
| 30% sucrose:L-Lysine (per mM) | 0.0856 | 0.0627 | 0.1096 | 8.9324 | 0.0004 |
| 30% sucrose:L-Lysine (at 10.0 mM) | 0.8556 | 0.6266 | 1.0958 | 135.2731 | 0.0004 |
| L-Valine (at 10.0 mM) | -0.0853 | -0.2400 | 0.0606 | -8.1725 | 0.1230 |
| 30% sucrose:L-Valine (at 10.0 mM) | 0.2030 | -0.0287 | 0.4374 | 22.5089 | 0.0428 |

**Table 7.** Summary statistics of *Hypothesis 4* (L-Lysine, sucrose and L-Valine) with AUC of the time-series growth curve as the dependent variable in different treatment conditions.

| Term | Estimate | CI 2.5% | CI 97.5% | % Change | p-value |
|---|---|---|---|---|---|
| Intercept | 10.7598 | 10.6911 | 10.8380 | | 0.0002 |
| Acetic acid | -1.4278 | -1.7564 | -1.1741 | -76.0175 | 0.0002 |
| L-Glutamine (per mM) | -0.0078 | -0.0193 | 0.0031 | -0.7770 | 0.0824 |
| L-Glutamine (at 10.0 mM) | -0.0780 | -0.1926 | 0.0309 | -7.5039 | 0.0824 |
| Acetic acid:L-Glutamine (per mM) | -0.0437 | -0.0784 | -0.0049 | -4.2788 | 0.0152 |
| Acetic acid:L-Glutamine (at 10.0 mM) | -0.4373 | -0.7840 | -0.0490 | -35.4223 | 0.0152 |
| L-Leucine (at 10.0 mM) | -0.9040 | -1.0583 | -0.7693 | -59.5059 | 0.0002 |
| Acetic acid:L-Leucine (at 10.0 mM) | -0.4604 | -0.8109 | -0.0443 | -36.8946 | 0.0166 |

**Table 8.** Summary statistics of *Hypothesis 5* (L-Glutamine, acetic acid and L-Leucine) with AUC of the time-series growth curve as the dependent variable in different treatment conditions.

| Term | Estimate | CI 2.5% | CI 97.5% | % Change | p-value |
|---|---|---|---|---|---|
| Intercept | 10.9362 | 10.8886 | 10.9845 | | 0.0002 |
| Lithium chloride | -1.8723 | -1.9975 | -1.7507 | -84.6234 | 0.0002 |
| L-Arginine (per mM) | -0.0417 | -0.0607 | -0.0232 | -4.0813 | 0.0002 |
| L-Arginine (at 5.0 mM) | -0.2083 | -0.3034 | -0.1159 | -18.8076 | 0.0002 |
| Lithium chloride:L-Arginine (per mM) | -0.0017 | -0.0377 | 0.0352 | -0.1678 | 0.4697 |
| Lithium chloride:L-Arginine (at 5.0 mM) | -0.0084 | -0.1887 | 0.1760 | -0.8360 | 0.4697 |
| L-Alanine (at 5.0 mM) | -0.1017 | -0.1818 | -0.0269 | -9.6668 | 0.0048 |
| Lithium chloride:L-Alanine (at 5.0 mM) | -0.1726 | -0.3571 | 0.0140 | -15.8520 | 0.0358 |

**Table 9.** Summary statistics of *Hypothesis 6* (L-Arginine, lithium chloride and L-Alanine) with AUC of the time-series growth curve as the dependent variable in different treatment conditions.

| Term | Estimate | CI 2.5% | CI 97.5% | % Change | p-value |
|---|---|---|---|---|---|
| Intercept | 10.6823 | 10.5761 | 10.7985 | | 0.0002 |
| (S)-lactic acid | -0.2747 | -0.4346 | -0.1291 | -24.0235 | 0.0006 |
| L-Proline (per mM) | -0.0064 | -0.0144 | 0.0016 | -0.6413 | 0.0584 |
| L-Proline (at 20.0 mM) | -0.1287 | -0.2885 | 0.0311 | -12.0745 | 0.0584 |
| (S)-lactic acid:L-Proline (per mM) | -0.0053 | -0.0170 | 0.0066 | -0.5290 | 0.1880 |
| (S)-lactic acid:L-Proline (at 20.0 mM) | -0.1061 | -0.3394 | 0.1326 | -10.0656 | 0.1880 |
| L-Alanine (at 20.0 mM) | -0.6429 | -0.3365 | -38.5959 | 0.0002 | |
| (S)-lactic acid:L-Alanine (at 20.0 mM) | -0.1883 | -0.4411 | 0.0615 | -17.1662 | 0.0684 |

**Table 10.** Summary statistics of *Hypothesis 7* (L-Proline, lactic acid and L-Alanine) with AUC of the time-series growth curve as the dependent variable in different treatment conditions.

| Term | Estimate | CI 2.5% | CI 97.5% | % Change | p-value |
|---|---|---|---|---|---|
| Intercept | 10.9446 | 10.8981 | 10.9874 | | 0.0002 |
| Formic acid | -2.0004 | -2.2581 | -1.7573 | -86.4722 | 0.0002 |
| Aminoadipate (per mM) | -0.2776 | -0.2958 | -0.2617 | -24.2421 | 0.0002 |
| Aminoadipate (at 5.0 mM) | -1.3881 | -1.4788 | -1.3085 | -75.0460 | 0.0002 |
| Formic acid:Aminoadipate (per mM) | 0.0768 | -0.0083 | 0.1610 | 7.9840 | 0.0388 |
| Formic acid:Aminoadipate (at 5.0 mM) | 0.3841 | -0.0415 | 0.8052 | 46.8243 | 0.0388 |
| L-Proline (at 5.0 mM) | 0.0011 | -0.0745 | 0.0770 | 0.1108 | 0.4789 |
| Formic acid:L-Proline (at 5.0 mM) | 0.0225 | -0.7586 | 0.6890 | 2.2737 | 0.4561 |

**Table 11.** Summary statistics of *Hypothesis 8* (aminoadipate, formic acid and L-Proline) with AUC of the time-series growth curve as the dependent variable in different treatment conditions.
